# Automatic classification of diatoms of Merja fouarate

**DOI:** 10.1101/099135

**Authors:** Nouzha Chahboune, Mohamed Mehdi, Allal Douira

## Abstract

**Objective:** This work consists of developing an automatic method of form analysis and classification of diatoms through processing of scanned images. The fundamental objective is to determine to what extent two diatoms are similar from the images stored in a database, and to conclude to what classes they belong. To do so, a comparison is made between a manual identification and an automatic identification, based on the ultimate points and the Freeman Chain Code.

## INTRODUCTION

We present in this study, an alternative to manual techniques of diatoms determination. The diatom identification and classification issue proves to be delicate, even for specialists. The number of known species is very large (approximately 11,000) and identification with the naked eye is a source of error (Kelly and al., 2002). These problems inevitably lead to a long, time-consuming and unreliable analysis (Guo, 2004; Loke, 2002). To address this problem, this study suggests the creation of an automated process based on techniques used in pattern recognition of diatoms, to identify and classify them automatically.

## MATERIAL AND METHODS

The automatic identification of diatoms relies on the software ‘ImageJ’; a digital image processing application developed at the National Institute of Health (Wayne, 1997).

### Development of a diatom image database

Before any identification, it is necessary to develop an image database. These are 10 species of diatoms from the Merja Fouarate, located in the plateau of the Mamora near the city of Kenitra in Morocco. This database is not a representative sample, since our goal here is to check the effectiveness of our method of identification. Subsequently, it would be possible to enhance the diatomic images database.

**Figure 3:**
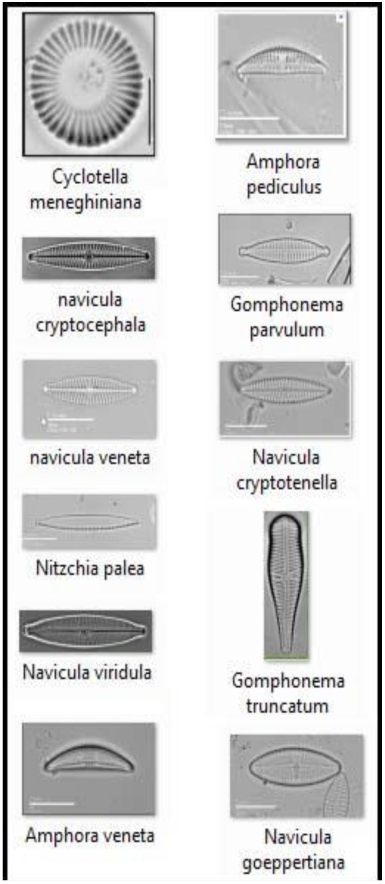
Example of the Merja fouarate diatoms.

The organization of the database is based on cross-indexing. The principle is: the attributes representing each diatom are stored in a vector. Each vector attribute is indexed to diatoms that have it and each diatom is indexed to the attributes which belong to it. During the identification, when an attribute is selected, the analysis continues from the diatoms related to this item, and in two or three iterations there remain only one or two candidates to identify.

### Diatom Scanning Method

The digital image processing based on mathematical morphology (Serra 1988) maps a picture that we try to identify with an image located in the image database (Bres S., and al. 2003). To enable this similarity search, the system retrieves the characteristics of the Diatom to be analyzed. These operations are to produce in matrix form, the outline and ornamentation. After these steps, it tries to find the corresponding diatom, according to the criteria that we developed above, namely the relevance of the items and the knowledge base.

### Outline Extraction

The classification of diatoms we do is based on the outline (Deriche, 2004). In fact this feature allows to organize the database of diatomic Images according to the species. This will reduce the number of candidates to analyze considerably. To do so, we proceed as follows: Process / Binary / Make Binary.

**Figure 4:**
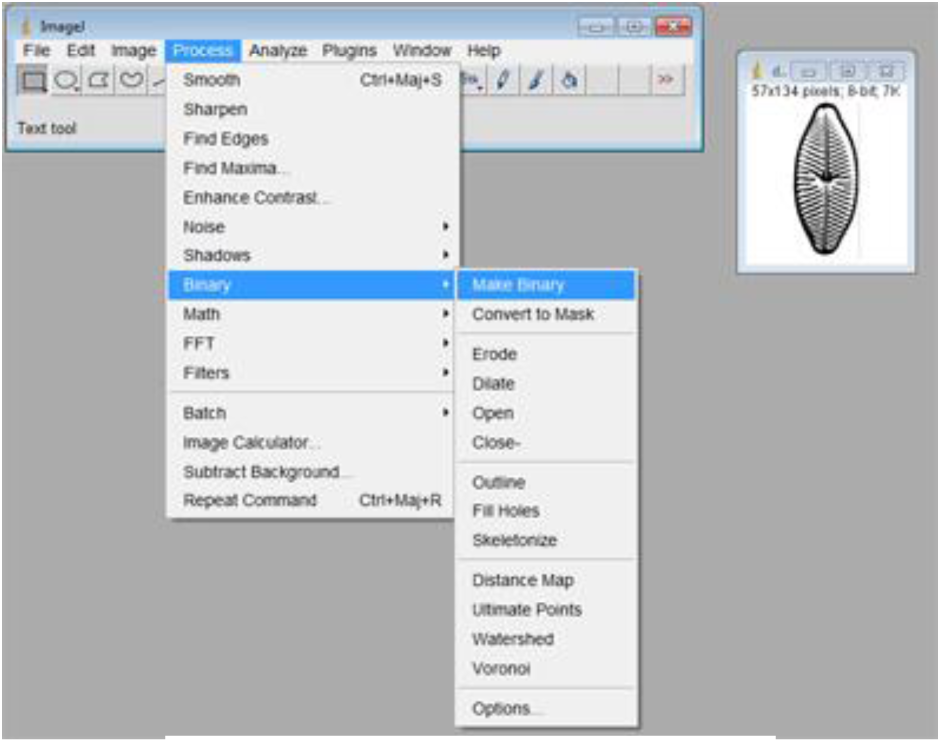
Binarization of the diatom.

Then we use the filling filter that darkens the interior, which can be achieved using the function *Process / Binary / Fill Holes.*

**Figure 5:**
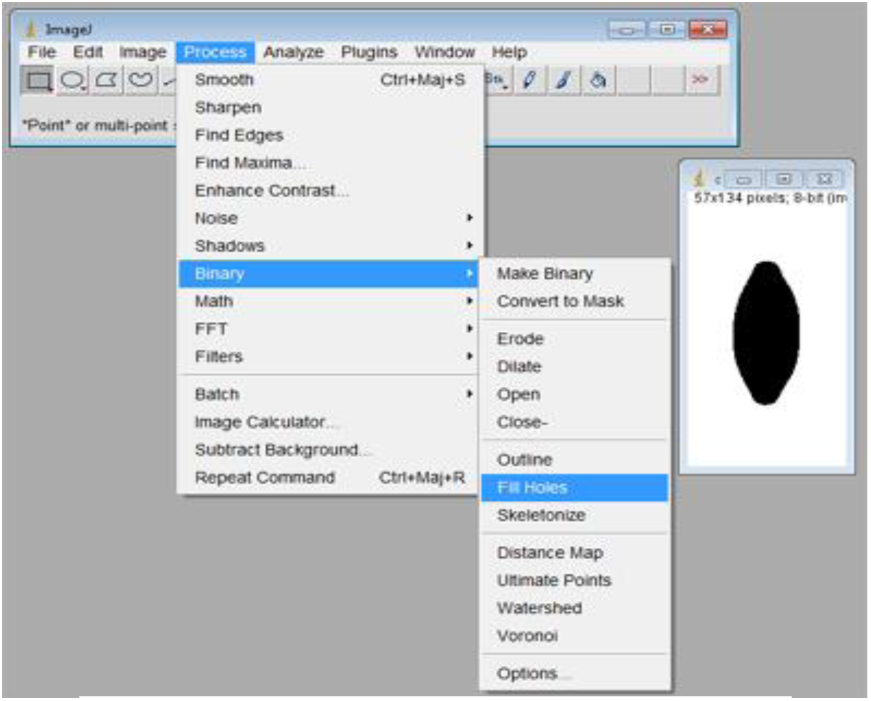
‘Fill Holes’ of the diatom image.

The third filter is based on the Sobel filter which allows to obtain the outline. The function that uses this filter is Process / Find Edges

**Figure 6:**
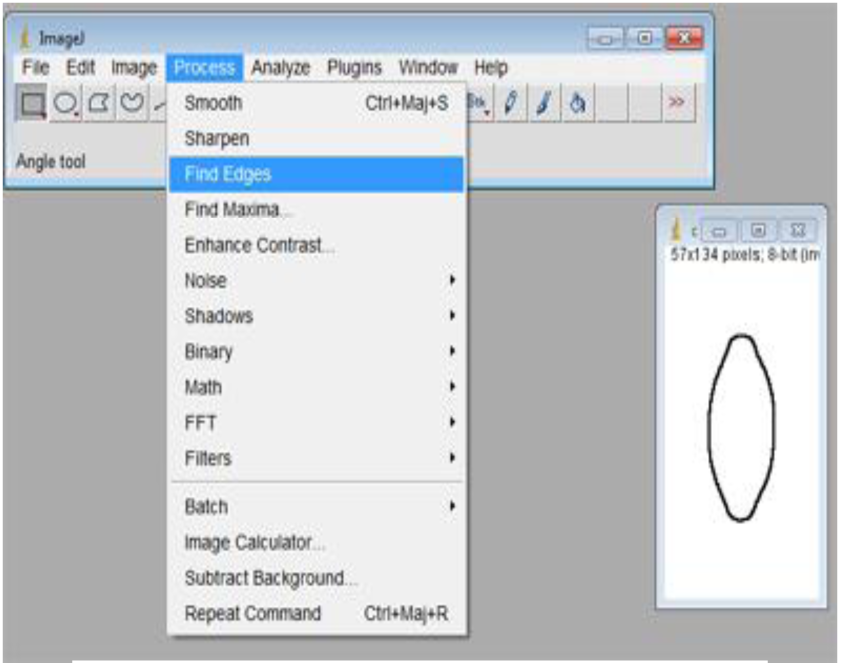
Extraction of the diatom outline.

The fourth filter is the skeletonization of the outline. Process / Binary / Skeletonize

**Figure 7:**
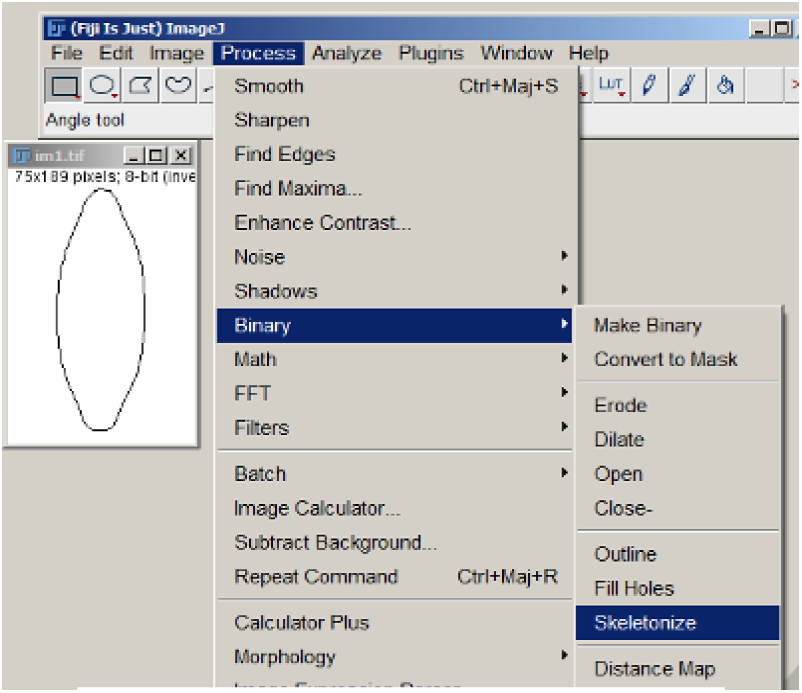
skeletonization of the diatom.

The fifth step is to obtain the corresponding vector by counting the number of even digits and the number of odd digits, according to the following formula: (741 x 1) + (151 × √2) ≈ 987.55

### Extraction of ornamentation

To recover the streaks of the pattern areas, we first perform a thresholding to obtain a binarized image. The image must be in grayscale (Dubuisson 1990).

**Figure 8:**
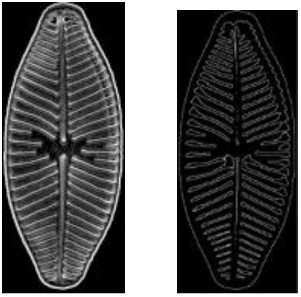
Binarization of the diatom image.

Upon completion of this step, we use a morphological operation that performs a subtraction between two matrices. In our case, it concerns the image that we have with the skeletonized outline image.

**Figure 9:**
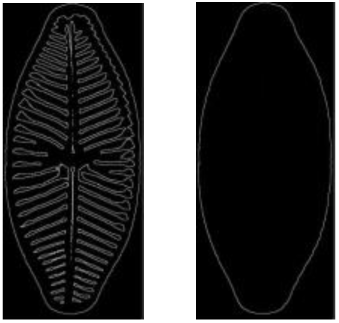
Determination of the outline of the diatom.

This allows us to get the frustule without the outline.

**Figure 10:**
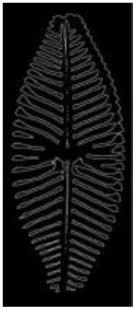
Determination of the background of the diatom.

This is done by using the command Image Calculator from the Menu Process, which calculates the difference between a binarized image and its dilated derivative, and features result in a third image.

**Figure 11:**
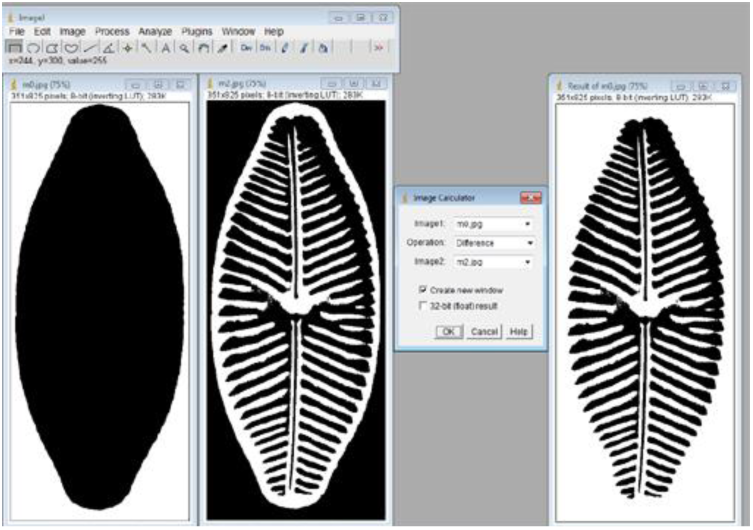
Extraction of the diatoms ornamentation.

## DETECTION OF ULTIMATE POINT ZONES

This step allows us to extract the characteristics of the candidate diatom. These characteristics correspond to ultimate points. We relied on the ultimate point detector to analyze the ornamentation. The ultimate points are characteristic points of the image that particularly hold information. They (ultimate points) have the property to hold single-handedly a summarized representation of an image, by synthesizing the information contained in the different pixels. The extraction of visual descriptors from the entire image allows the reduction of the required calculations, the database volume as well as the cost of the research of the most similar images. This allows not taking in consideration every pixel of the attributes while analyzing, but instead to sample only a representative subset. Thus, a very small number of items in the range of a 100 can hold the most important information on the image, to be compared with the 100.000 pixels of an image corresponding to a diatom.

From a binarized and Skeletonize image of a diatom we extract the outline to keep nothing but the frustule then we applies the ultimate point detection function on the frustule. To do this, on the ImageJ menu, we select the « Process » command, then « Binary » and finally « Ultimate Points », which is the function that produces the ultimate points.

**Figure 62:**
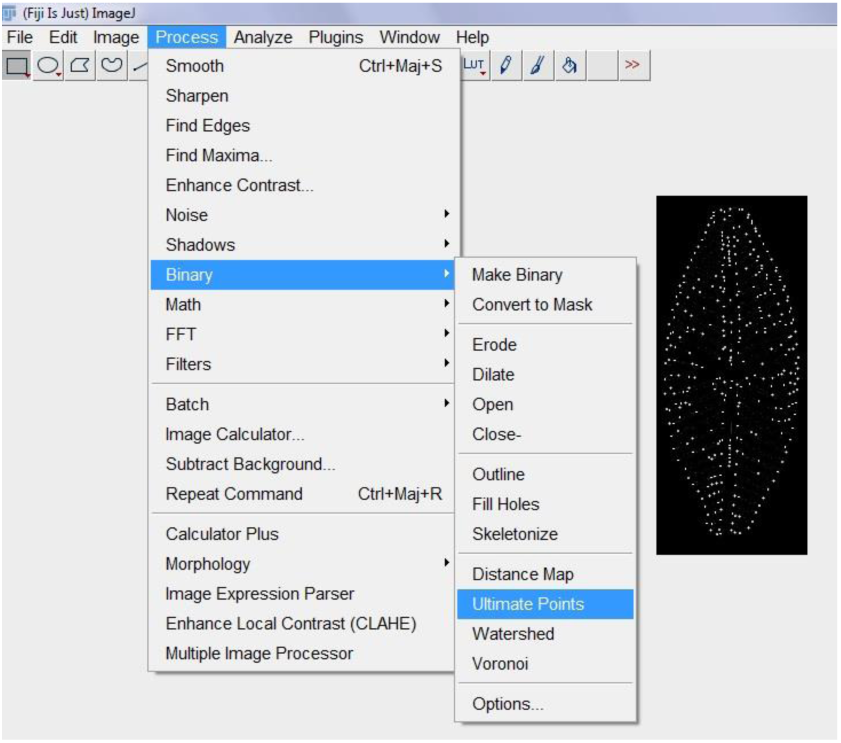
Ultimate points determination method on ImageJ.

## METHODOLOGY AND RESULTS

### Automatic identification of diatoms

by comparing two similar diatoms, we will show how automatic identification could solve the aforementioned problems.

We saw, through material and methods that the diatoms **are represented in the image database by their shape and their ornamentation in the form of a vector. These are the two criteria on which we will base our comparisons to show the efficiency of the automatic identification, Note that each vector contains the name of the diatom and its chain code (Freeman, 1961) determining the outline and the list of points corresponding to the ultimate points.**

### First step

Assume we want to identify the following diatom of which we do not know the identity:

**Figure 12:**
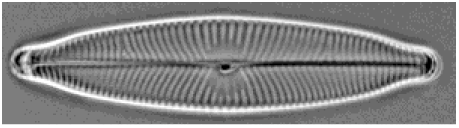
Targeted diatom.

An ImageJ macro analyses the image, which allow us to obtain the following characteristics:

**Figure 13:**
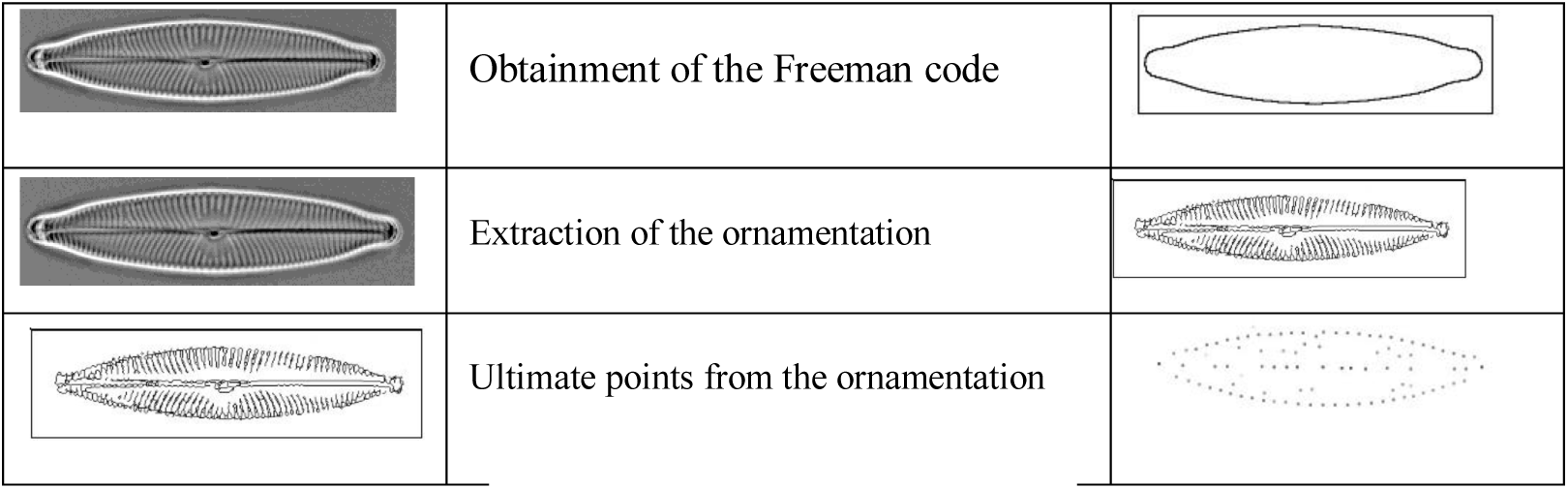
characteristics extraction.

### Second step

The first characteristic, allows us to obtain the Freeman code. Knowing that we obtain a code from each outline, in this case we only compare the codes to one another.

According to our example, an approximate comparison from the code gives us the following results:

**Figure 14:**
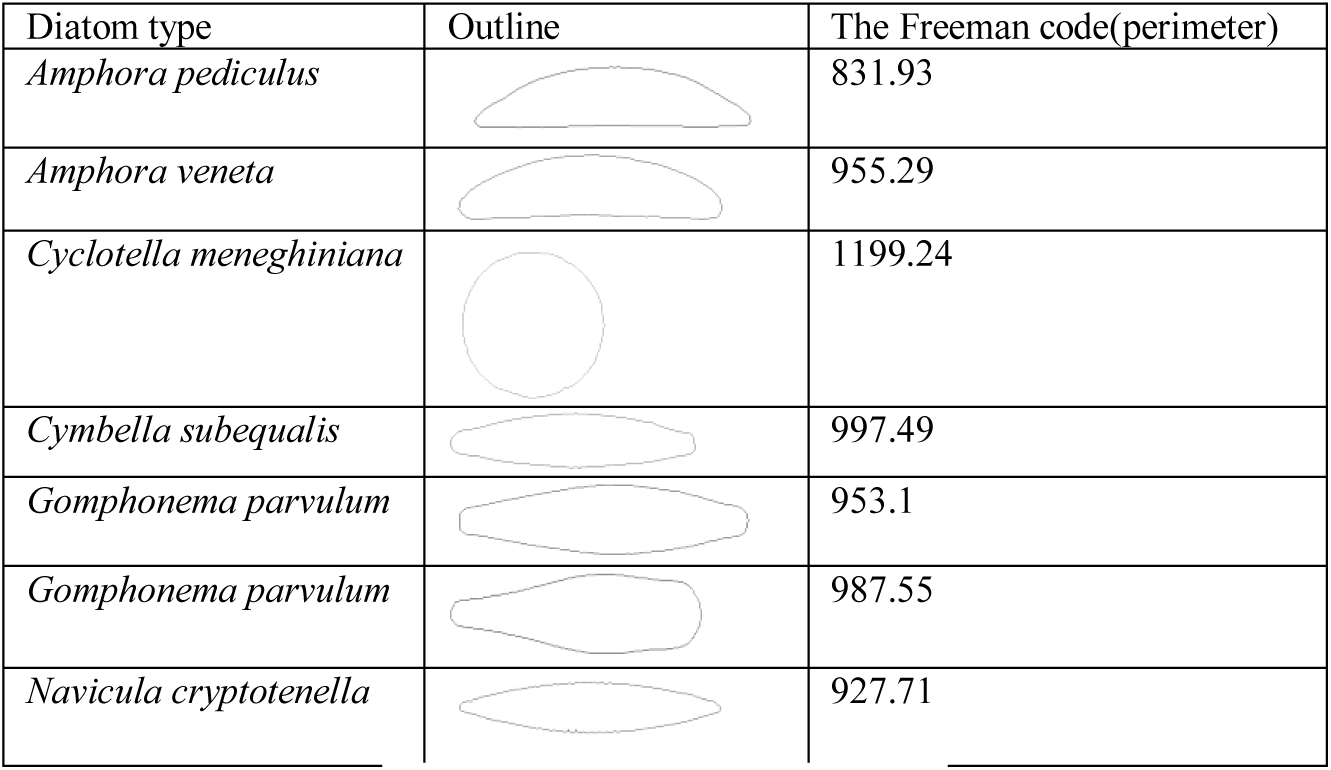
Freeman codes results.

### Third step

the second characteristic allows to compare the candidate image ornamentation to the targeted image, to do so we perform a comparison based on the ultimate points. We start by scaling down the image, we choose a 32/32 size, which allows us to compress the image without losing information. These ultimate points are represented as a vector.

**Figure 15:**
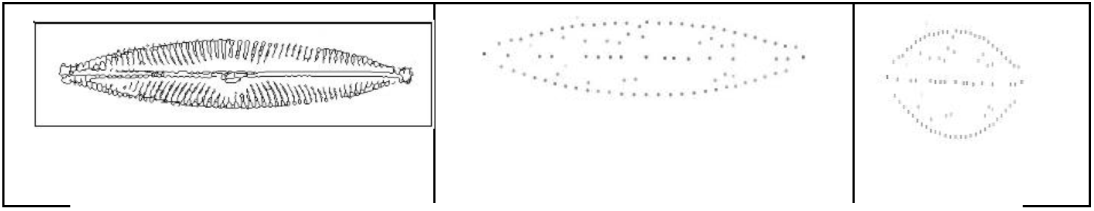
representation of the diatom’s ultimate points.

Thus, the outline allows selection of a set of diatoms, then we compare using cross-indexing, the vectors corresponding to the ultimate points of this set.

**Figure 16:**
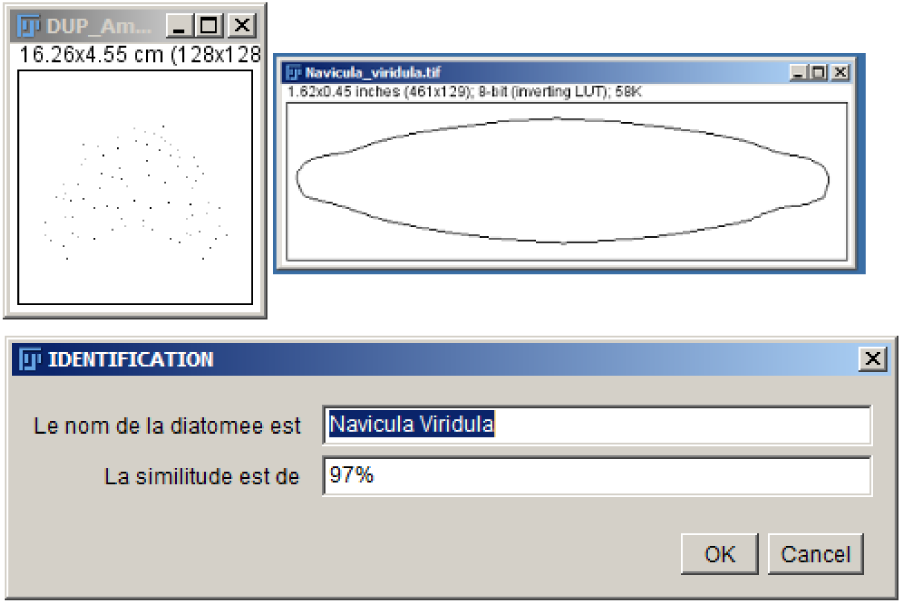
The diatom’s automatic identification result.

In our example, the last remaining candidate is *«Navicula Viridula» which matches our targeted image.*

## CONCLUSION AND RESEARCH APPLICATION

The world of diatoms is diversified into several groups, genera and species. It presents many morphological variations which are basic elements as to their determination and classification. Automatic image recognition provides valuable assistance for both the determination of diatoms, and as a teaching tool to assist identification to the benefit of new diatomists.

Our study aimed to automate the diatom identification process based on analysis and image interpretation techniques. Indeed, the image classification requires a candidate image, the one that must be identified and an image database in which to find the corresponding image, allowing identification. Comparing images is problematic. The first problem is the choice of the classification resolution. Indeed, the analysis of raw images is consuming in calculation time. It is therefore interesting to only compare what is necessary. Thus we have not taken into consideration all the attributes’ pixels to analyze, but we have taken a representative subset, based on the approach of “ultimate points”. The second problem concerns the classification method, this phase is particularly challenging because the slightest difference between the two characteristics sets can fail the identification. To avoid this issue, we recommend the use of a method based on approximate similarity. The use of a similarity coefficient has allowed us to have a little more flexibility against an overly discriminating binary comparison

Upon completion of this project, the results were very satisfactory, giving correct classification rates higher than 96% using all the methods discussed and even surpassing the results of human experts. However, the specific nature of certain potentially more complex species could not be resolved. Nevertheless research continues and relies heavily on the outcomes of Artificial Intelligence.

